# Dual therapy with corticosteroid ablates the beneficial effect of DP2 antagonism in chronic experimental asthma

**DOI:** 10.1101/2023.11.12.566772

**Authors:** Md Ashik Ullah, Sonja Rittchen, Jia Li, Bodie F. Curren, Muhammed Mahfuzur Rahman, Md Al Amin Sikder, Ridwan B. Rashid, Natasha Collinson, Mary Lor, Mark L. Smythe, Simon Phipps

## Abstract

**Background:** Prostaglandin D2 (PGD2) signals via the DP1 and DP2 receptors. In Phase II trials, DP2 antagonism decreased airway inflammation and airway smooth muscle (ASM) area in patients with moderate-to-severe asthma, but in the Phase III clinical trials, DP2 antagonism failed to significantly lower the rate of exacerbations. Here, we hypothesised that DP2 antagonism resolves established ASM remodeling via endogenous PGD2/DP1 activation and that this beneficial effect is ablated by dual corticosteroid therapy.

**Methods:** Neonatal mice were co-exposed to pneumonia virus of mice (PVM) and cockroach extract in early life to induce severe bronchiolitis, then re-infected with PVM and challenged to cockroach extract in adulthood to progress disease to chronic experimental asthma (CEA). The efficacy of DP2 antagonism monotherapy or various dual therapies was assessed in the setting of a rhinovirus (RV)-induced exacerbation.

**Results:** RV inoculation increased PGD2 release, mucus production, collagen deposition, transforming growth factor (TGF)-β1 expression and type-2 inflammation. Treatment with a DP2 antagonist or DP1 agonist ablated the aforementioned phenotypes, increased type-1 immunity, and decreased ASM area. Dual DP1-DP2 antagonism or dual corticosteroid/DP2 antagonism, which attenuated endogenous PGD2 levels, prevented the resolution of ASM area induced by DP2 antagonism alone. The resolution of ASM remodelling following DP2 antagonism was mediated by IFN-γ and associated with decreased TGF-β1 expression.

**Conclusion:** DP2 antagonism resolved ASM remodelling via PGD2/DP1-mediated upregulation of interferon-γ expression. Dual DP2 antagonism/corticosteroid therapy, as occurred in many of the human trials, suppressed PGD2 and IFN-γ production, impairing the efficacy of DP2 antagonism.

## INTRODUCTION

Acute exacerbation of chronic asthma, most commonly as a result of rhinovirus (RV) infection, causes significant morbidity, mortality and healthcare costs.^1^ A critical pathological feature of chronic asthma is airway remodelling characterized by goblet cell hyperplasia, airway smooth muscle (ASM) hyperplasia, and reticular basement membrane thickening. In an acute exacerbation of chronic asthma, the remodelled airways undergo bronchoconstriction causing airway narrowing and air-trapping leading to increased wheezing and breathlessness.^2^ Commonly used treatments, including corticosteroids or non-steroidal anti-inflammatory drugs, fail to halt or reverse ASM remodelling,^3^ while the emerging monoclonal antibody therapies have not been assessed sufficiently,^4^ and hence the prevailing view remains that ASM remodelling, once established, is irreversible. Although bronchial thermoplasty reduces ASM mass, this procedure is only recommended for individuals with uncontrolled-severe asthma, which limits its therapeutic application.^5^ Thus, there is an unmet need for new therapeutics to reverse established airway remodelling.

In addition to the type-2 ‘instructive cytokines’ thymic stromal lymphopoietin, IL-33, and IL-4, the eicosanoid prostaglandin D2 (PGD2) has been shown to amplify type-2 inflammation by activating multiple effector cells, including CD4^+^ T helper 2 cells, type 2 innate lymphoid cells (ILC2), eosinophils, mast cells and basophils.^6^ Additionally, PGD2 directly activates ASM cells and is a potent bronchoconstrictor.^7^ These effects are mediated via activation of the PGD2 receptor 2 (DP2), and this PGD2-DP2 pathway is associated with severe, poorly controlled, type-2-high asthma.^8^ Consequently, several DP2 antagonists were developed for the treatment of asthma, and in phase-II clinical trials, DP2 blockade was shown to decrease airway eosinophilia, improve lung function and delay recurrence of an asthma exacerbation in patients with mild-to-moderate asthma or inadequately controlled, severe asthma. ^9–15^ Intriguingly, in patients with persistent eosinophilic asthma, DP2 antagonism was observed to lower ASM area,^11^ although the mechanism of action remains unknown. Despite these promising findings, large-scale phase III trials in patients with uncontrolled or severe asthma found that the DP2 antagonist fevipiprant only modestly (and not significantly) decreased the rate of exacerbations compared with placebo, and failed to improve lung function.^16,17^ Similarly, in a Phase IIb study, GB001 failed to significantly reduce the odds of asthma worsening.^18^ Precisely why the DP2 antagonists were ineffective in these latter trials remains unclear, as does the mechanism of action by which fevipiprant successfully reverses ASM remodelling.

In a pre-clinical model of viral bronchiolitis and using primary human bronchial epithelial cells, we previously demonstrated that the beneficial effects of DP2 antagonism, including the attenuation of type-2 inflammation and the restoration of antiviral immunity, are largely mediated via the activation of the PGD2 receptor DP1,^19^ indicating that the beneficial effects of DP2 antagonism are dependent on the production of endogenous PGD2. As corticosteroids inhibit and downregulate the expression of cyclooxygenase (COX)2 and subsequent PGD2 production, we hypothesized that the continuation of standard-of-care corticosteroid treatment, as occurred in the majority of trials, may have decreased the effectiveness of DP2 antagonismin in the treatment of asthma. Here, using an established high-fidelity pre-clinical mouse model of chronic experimental asthma (CEA), characterized by eosinophilic inflammation and persistent ASM remodelling,^20–22^ we sought to test this hypothesis and to investigate the molecular mechanism by which DP2 antagonism reverses ASM remodelling.

## METHODS

Additional details are described in the supplementary section (Appendix S1). The details of all the reagents used are listed in Supplementary Table S1.

### Rhinovirus-induced exacerbation model

Neonatal BALB/c mice were intranasally (i.n. route) inoculated with pneumonia virus of mice (PVM; 1 PFU) at postnatal day (PND) 7 and cockroach extract (CRE, 1 µg; i.n. route) at 3 days post-infection (dpi). Mice were challenged with PVM (100 PFU) at 42 dpi and CRE (1 μg) at 45, 52, 59 and 66 dpi to induce chronic experimental asthma (CEA).^20^ Four weeks later, mice were inoculated with vehicle or RV-1b (5×10^6^ median tissue culture infective dose [TCID50]) to induce an acute exacerbation.^21–23^ Where indicated, separate groups of mice were treated with diluent: a DP1-specific agonist, BW245c (1 mg/kg of body weight; i.n.); a DP1-specific antagonist, MK-0524 (5 mg/kg of body weight; i.n.), a DP2-specific antagonist, timapiprant also known as OC000459 (10 mg/kg of body weight; oral gavage), or a hematopoietic prostaglandin D synthase (h-PGDS) inhibitor, PK007, generated in the laboratory of Dr. Mark Smythe (10 mg/kg of body weight; oral gavage) from postnatal day 99 until euthanasia. In some experiments, mice were treated with soluble IL-13Rα2 (i.n., 200 µg/dose; Pfizer, NY, USA) on alternate days starting from postnatal day 100 or fluticasone (i.n., 1 mg/kg of body weight; Sigma-Aldrich, MO, USA) daily from postnatal day 99 onwards, and the mice euthanized at 1, 3, and 7 dpi. Where indicated, anti-IFN-γ (8 mg/kg of body weight) antibody was administered intraperitoneally on alternate days starting at postnatal day 100. All studies were approved by the Animal Ethics Committee of QIMR Berghofer Medical Research Institute. Mice were euthanized at the indicated time points, and different tissue samples were harvested and processed for immunological assays as outlined in the supplementary methods (Appendix S1).

## RESULTS

### Treatment with a DP2 antagonist, but not a corticosteroid, ameliorates airway remodelling during a RV-triggered exacerbation of chronic experimental asthma

To investigate the immune mechanisms by which DP2 antagonism reverses established airway remodelling, we employed a high-fidelity mouse model of CEA previously established in our laboratory.^20,21^ Inoculation of WT mice from infancy with the respiratory virus PVM (a close relative of human respiratory syncytial virus) and cockroach extract (CRE) (Figure 1A), leads to the development of CEA, as demonstrated by persistent airway smooth muscle (ASM) remodelling and collagen deposition.^20,21^ The model simulates the human epidemiology linking severe/frequent lower respiratory infections and allergic sensitization that synergistically increase the risk of childhood asthma.^24^ To mimic an acute asthma exacerbation, mice were inoculated with RV four weeks later, and immunopathology was assessed at 1, 3, and 7 days post-infection (dpi). Consistent with our previous findings and that of others, ^19,25,26^ respiratory viral infection led to a significant increase in PGD2 levels in the lung compared to non-infected control mice (Figure 1B), suggesting that PGD2 contributes to viral-triggered exacerbations of asthma. Treatment with a DP2 antagonist (timapiprant, also known as OC000459) or a DP1 agonist (BW245c) prior and during the RV infection (Figure 1A) had no effect on lung PGD2 levels at 1 or 7 dpi, although DP2 antagonism increased PGD2 levels at 3 dpi (Figure 1B). Consistent with our previous findings, ASM area and peri-bronchial collagen deposition were significantly elevated in mice with CEA compared to naïve controls (Figure 1C-F).^22^ Neither pathology was further increased following RV exposure; however, treatment with the DP2 antagonist during RV infection significantly decreased ASM area, mirroring the findings of the phase II clinical trial of fevipiprant^11^ and decreased peri-bronchial collagen (Figure 1C, D and F). A similar outcome was observed following treatment with a DP1 agonist (Figure 1C, D and F). We compared this treatment effect to fluticasone, a mainstay treatment for asthma, and to soluble (s)IL-13Rα2, a treatment that neutralizes IL-13, a central mediator of type-2 inflammation. Consistent with clinical investigations,^27,28^ fluticasone had little effect on ASM area or collagen deposition. sIL-13Rα2 similarly had no effect ASM area but did lower collagen deposition (Figure 1E and G). RV inoculation increased mucus production at 3 and 7 dpi (Figure 1H), and this effect was attenuated by DP2 antagonism or DP1 agonism as well as by fluticasone or sIL-13Rα2 treatment (Figure 1H-I). Collectively, these data indicate that DP2 antagonism or DP1 agonism during a virus-induced exacerbation of asthma resolves established airway remodelling, including the increased ASM mass, whereas corticosteroid treatment only prevents the increase in mucus production.

**FIGURE 1.**
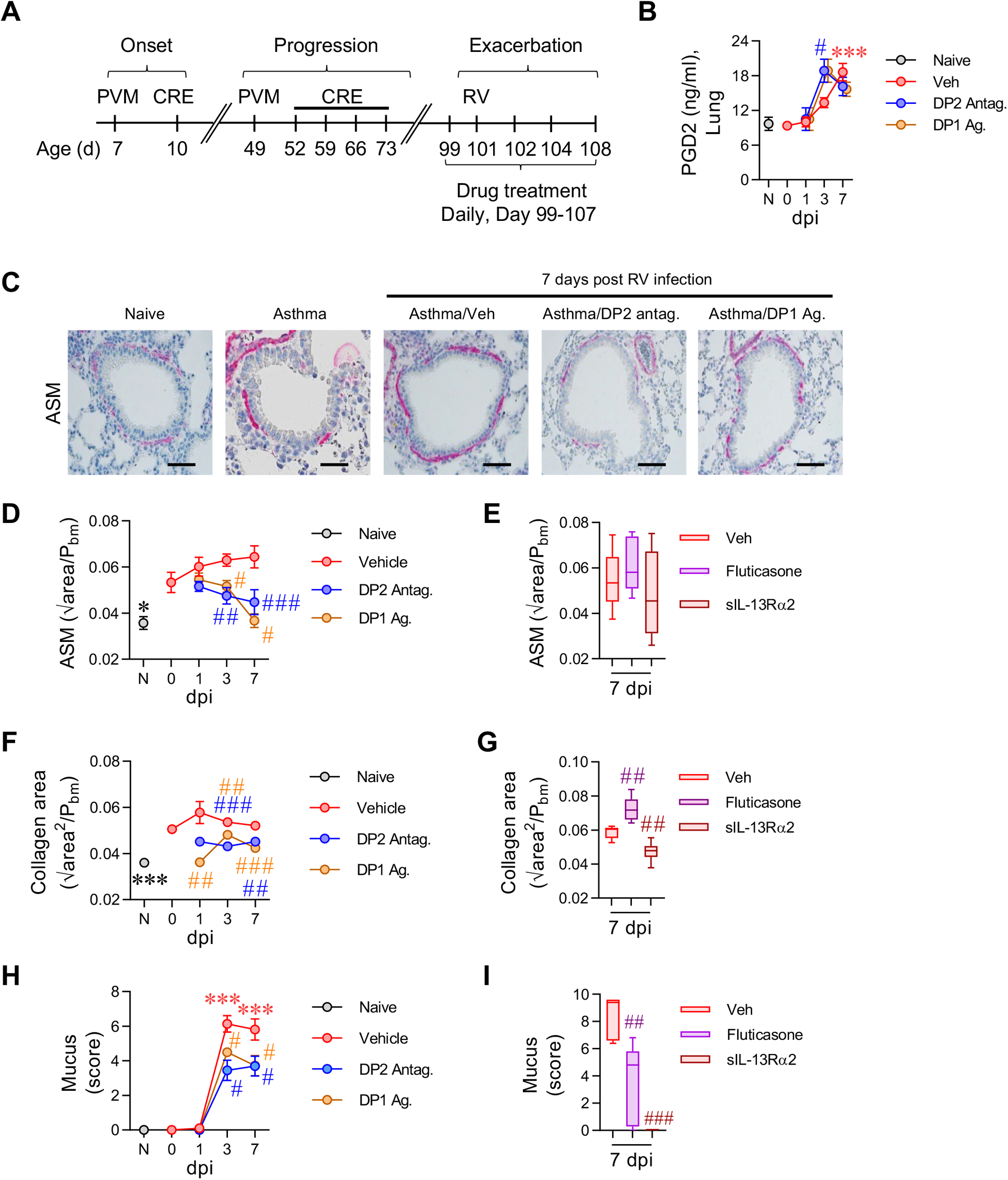
Treatment with a DP2 antagonist, but not a corticosteroid, ameliorates airway remodelling during a RV-triggered exacerbation of CEA. (A) Study design. Seven days old mice were inoculated with PVM and exposed to CRE 3 days later. Mice were re-infected with PVM six weeks later and exposed to CRE weekly for four weeks. Four weeks later, mice were inoculated with RV-1b. Separate groups of mice were treated with a DP2 antagonist (OC000459), DP1 agonist (BW245c), fluticasone, soluble IL-13Rα2 or vehicle. Mice exposed to vehicle instead of virus or allergen were referred to as naïve. Mice were then inoculated with RV-1b and euthanized at 1, 3 and 7 days post infection (dpi). (B) Lung PGD2 levels. (C) Representative lung histology of α-smooth muscle actin (SMA) expression. Scale bars = 50µm. (D-E) Airway smooth muscle (ASM) area. (F-G) Collagen deposition (H-I) Mucus score. Data are presented as mean ± SEM or box-and-whisker plots, and are pooled data from two independent experiments (n = 4-16 mice per group). Statistical significance between different time points or different groups was determined using one-way ANOVA with Dunnett’s multiple comparison test. * denotes p<0.05, ** denotes p<0.01 and *** denotes p<0.001 compared to Vehicle group. # denotes p<0.05, ## denotes p<0.01 and ### denotes p<0.001 compared to RV-infected group at corresponding time point.

### DP2 antagonism or DP1 agonism decreases lung TGF-β1 expression

Transforming growth factor β1 (TGF-β1) contributes to ASM remodelling and collagen synthesis and is elevated in the lungs of patients with asthma.^29,30^ Consistent with this, we observed increased TGF-β1 immunoreactivity around the airways of mice with CEA compared to their naïve counterparts (Figure 2A-C). Following RV inoculation, TGF-β1 expression was further increased around the airways, a phenotype confirmed by measuring TGF-β1 (by ELISA) in whole lung homogenates (Figure 2A-B, D). Significantly, DP2 antagonism or DP1 agonism ablated this increase in TGF-β1 expression and lowered TGF-β1 levels to those observed in naïve controls (Figure 2A-B, D). In contrast, neither fluticasone nor sIL-13Rα2 treatment affected TGF-β1 expression (Figure 2C and E). These data suggest that DP2 antagonism or DP1 agonism resolves airway remodelling by suppressing the expression of TGF-β1.

**FIGURE 2.**
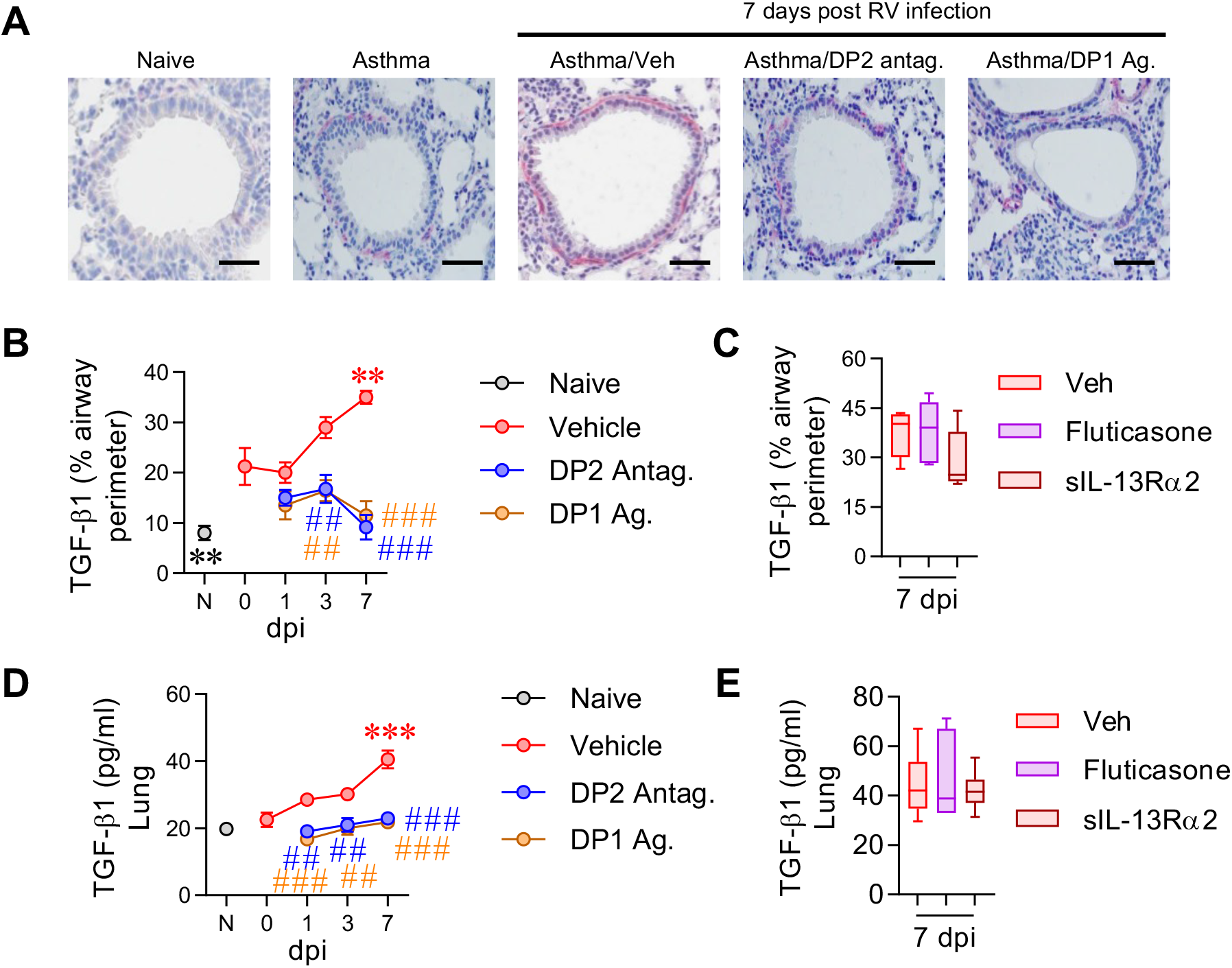
DP2 antagonism or DP1 agonism decreases lung TGF-β1 expression. (A) TGF-β1 immunoreactivity (red colouration). Scale bars = 50µm. (B-C) Quantification of TGF-β1 immunoreactivity around the airways. (D-E) Lung TGF-β1 levels in homogenates. Data are presented as mean ± SEM or box-and-whisker plots and are pooled data from two independent experiments (n = 4-9 mice per group). Statistical significance between different time points or different groups was determined using one-way ANOVA with Dunnett’s multiple comparison test. ** denotes p<0.01 and *** denotes p<0.001 compared to vehicle group. ## denotes p<0.01 and ### denotes p<0.001 compared to RV-infected group at corresponding time point.

### DP2 antagonist-mediated resolution of airway remodelling is dependent on endogenous PGD2 and ablated by dual therapy with a corticosteroid

The effectiveness of the DP1 agonist in resolving established ASM remodelling suggested that DP2 antagonism primarily elicits its beneficial effects via DP1 receptor activation, and thus requires endogenous PGD2. To test this, we hypothesized that the beneficial effects of DP2 antagonism (i) would not occur in the absence of RV-induced PGD2 release, (ii) would be ablated by dual DP1/DP2 antagonism, and (iii) would not be replicated by inhibition of hematopoietic prostaglandin D synthase (h-PGDS), since this lowers endogenous PGD2, ablating the activation of both DP1 and DP2. As predicted, treatment with a DP2 antagonist in the absence of a RV-induced exacerbation (akin to ‘stable’ asthma) failed to affect ASM area, collagen deposition, or TGF-β1 expression (Figure S1). Notably, in RV-inoculated mice with CEA, dual antagonism with a DP2 antagonist and a DP1-specific antagonist (MK0524) prevented the resolution of airway remodelling and attenuated the fall in TGF-β expression mediated by DP2 antagonism (Figure 3A-D).

**FIGURE 3.**
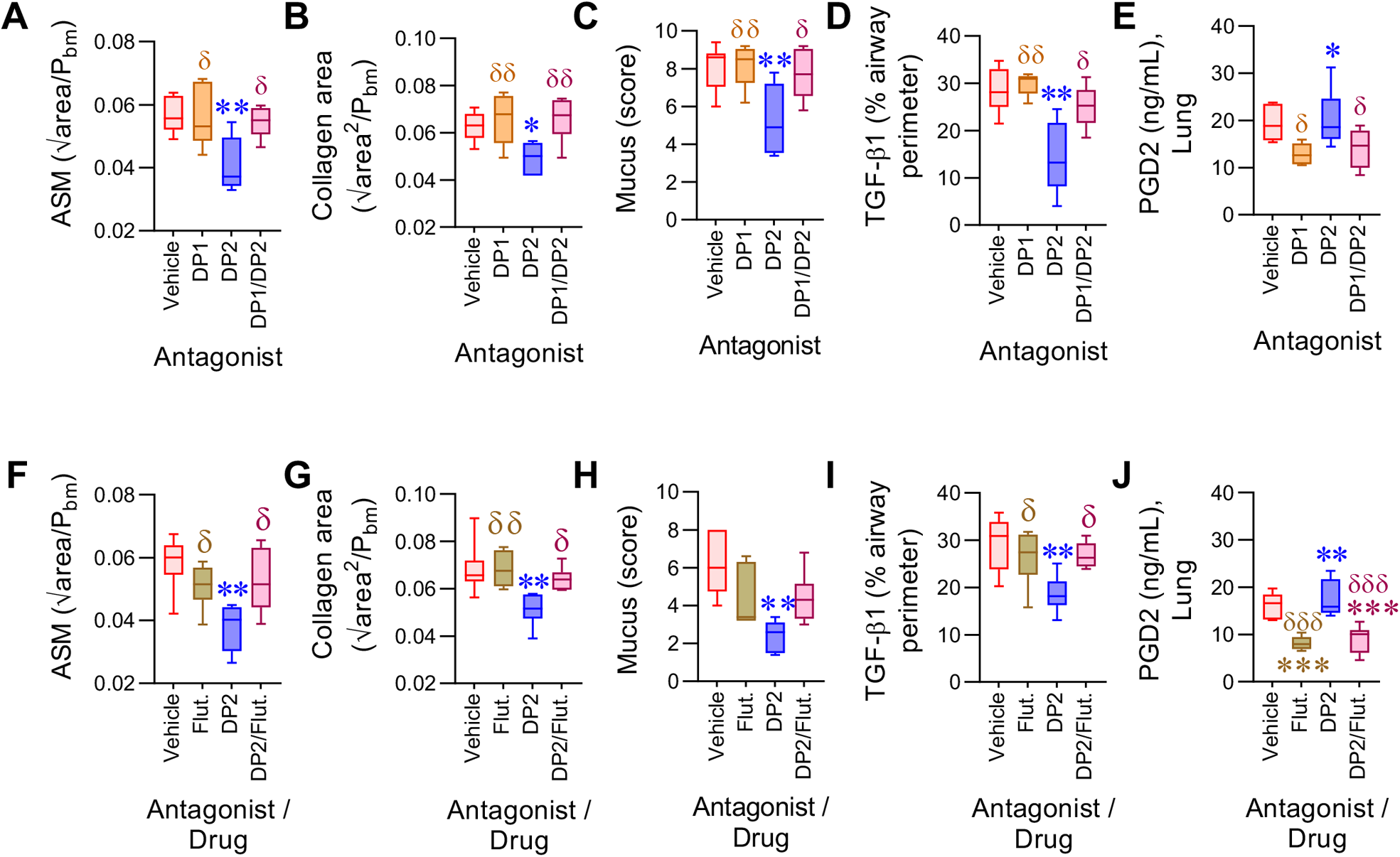
DP2-antagonist mediated resolution of airway remodelling is dependent on endogenous PGD2 and ablated by dual therapy with a corticosteroid. (A-E) Mice with CEA were Inoculated with RV-1b and given vehicle, DP2 antagonist or DP1 antagonist or both DP2 and DP1 antagonist daily from day 99 and euthanized at 7 dpi. Quantification of (A) ASM, (B) collagen deposition, (C) Mucus score and (D) TGF-β1 expression. (E) Lung PGD2 levels. (F-J) Mice with chronic asthma were infected with RV-1b and given vehicle, DP2 antagonist fluticasone or both DP2 antagonist and fluticasone daily from day 99 and euthanized at 7 dpi. Quantification of (F) ASM, (G) collagen deposition, (H) Mucus score and (I) TGF-β1 expression. (J) Lung PGD2 levels. Data are presented as box-and-whisker plots and are representative of two independent experiments (n = 5-7 mice per group). Statistical significance between different groups was determined using one-way ANOVA with Dunnett’s multiple comparison test. * denotes p<0.05, ** denotes p<0.01 and *** denotes p<0.001 compared to vehicle group. δ denotes p<0.05 and δδ denotes p<0.01 compared to DP2 antagonist treated group.

The dual DP1/DP2 antagonism lowered PGD2 production (Figure 3E), consistent with the notion that PGD2/DP1 activation acts in a positive feed-forward loop, which presumably enhances its efficacy. Despite decreasing PGD2 levels, h-PGDS inhibition failed to affect ASM area, collagen deposition, mucus hypersecretion or TGF-β1 expression (Figure S2A-E). The requirement for endogenous PGD2 suggested that the effectiveness of DP2 antagonism would be decreased by co-treatment with a corticosteroid, as corticosteroids inhibit cyclooxygenases and h-PGDS, lowering prostanoid production.^31,32^ Significantly, dual fluticasone/DP2 antagonism treatment ablated the beneficial effect of DP2 antagonism (Figure 3F-I), an effect associated with a significant decrease in lung PGD2 levels (Figure 3J).

### DP2 antagonism or DP1 agonism attenuates RV-induced type-2 inflammation and promotes type-1 immunity

Next, to explore the mechanism by which DP1 agonism reverses airway remodelling, we investigated the effect of DP2 antagonism or DP1 agonism on RV-induced airway inflammation (gating strategies in Figure S3). In response to RV, neutrophil numbers increased sharply at 1 dpi and then waned by 3 dpi, whereas eosinophil numbers rose steadily and peaked at 7 dpi (Figure 4A). Treatment with the DP2 antagonist or DP1 agonist decreased the infiltration of both granulocytes at 1 and 3 dpi, although at 7 dpi, both interventions led to an increase in lung neutrophils (Figure 4A-B). Similar to the temporal pattern of eosinophilia, GATA3^+^ T_H_2 cells and ILC2 numbers tended to increase between 3 and 7 dpi, and both cell types were decreased in response to DP2 antagonism or DP1 agonism (Figure 4B). The type-2 effector cytokines, IL-4, IL-5, and IL-13 were elevated, particularly at 1 and 3 dpi, and significantly decreased by either treatment (Figure 4C and Figure S4A). Similarly, IL-17A expression, which was elevated at 1 dpi, was ablated by DP2 antagonism or DP1 agonism at 1 dpi (Figure 4D) and aligned with a reduction in lung RORγt^+^ T_H_17 cells but not ILC3s (Figure S4B-C). In contrast, IFN-γ expression in mice with CEA was numerically (but not statistically) lower than that of naïve controls at baseline and decreased further at 3 and 7 dpi (Figure 4E). The pattern of IFN-γ expression was not associated with T-bet^+^ T_H_1 cells, ILC1s, or NK cells (Figure S4B-C). Following treatment with the DP2 antagonist or DP1 agonist, the expression of IFN-γ and TNF-α, another type-1 cytokine, was significantly increased (Figure 4E). Of note, dual DP1/DP2 antagonist or dual fluticasone/DP2 antagonist treatment abrogated the increase in IFN-γ and TNF-α expression induced by DP2 antagonist therapy alone (Figure 4F-G). Dual fluticasone/DP2 antagonist treatment also diminished the suppressive effect of DP2 antagonism monotherapy on IL-4 and IL-5 levels, although IL-13 levels remained significantly lower than the vehicle-treated mice (Figure S4D). Collectively, these data demonstrate that (i) DP2 antagonism or DP1 agonism attenuates type-2 inflammation and promotes type-1 immunity and (ii) dual DP2 antagonism/corticosteroid therapy prevents this shift to type-1 immunity.

**FIGURE 4.**
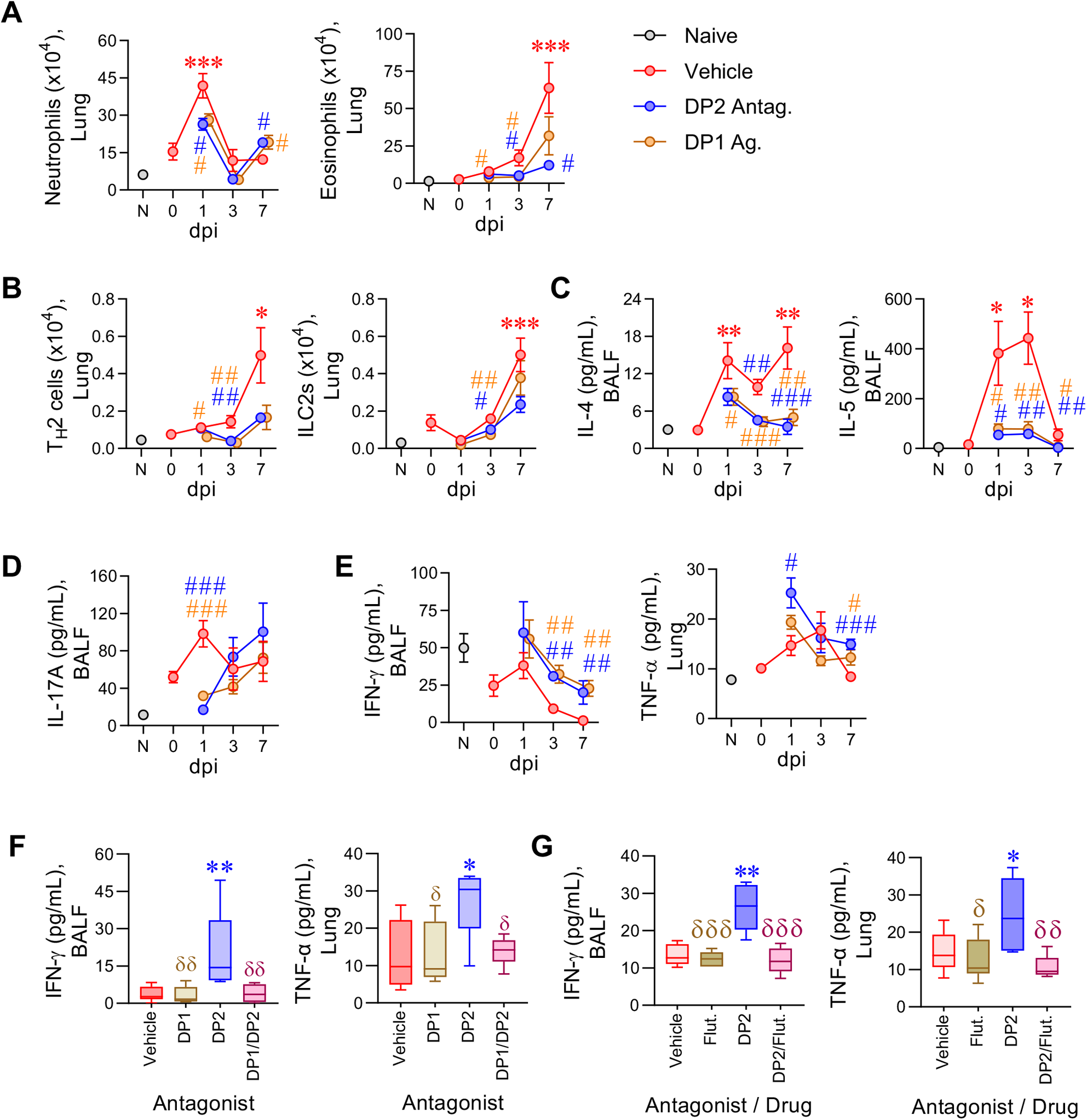
DP2 antagonism or DP1 agonism attenuates RV-induced type-2 inflammation and promotes type-1 immunity. Mice with CEA were inoculated with RV-1b and treated daily with a DP2 antagonist, a DP1 agonist, or vehicle starting from day 99. Mice were euthanized at 1, 3 and 7 dpi. (A) Number of neutrophils (SigF^−^CD11b^+^Ly6G^+^CD11c^−^) and eosinophils (SigF^+^CD11b^+^Ly6G^−^CD11c^−^) in the lungs. (B) Number of T_H_2 cells (GATA3^+^CD4^+^T) and ILC2s (CD3ε^−^CD19^−^CD45R^−^CD11c^−^Gr 1^−^CD11b^−^NK1.1^−^ CD90.2^+^CD200R1^+^ GATA3^+^) in the lungs. (C) Concentrations of IL-4 and IL-5 in the bronchoalveolar lavage fluid (BALF). (D) IL-17A expression in the BALF. (E) IFN-γ expression in the BALF and TNF-α expression in the lung. Data are presented as mean ± SEM and are representative of two independent experiments showing similar results (n = 4-11 mice per group). Statistical significance between different time points or different groups was determined using one-way ANOVA with Dunnett’s multiple comparison test. * denotes p<0.05; ** denotes p<0.01 and *** denotes p<0.001 compared to Vehicle group. # denotes p<0.05, ## denotes p<0.01 and ### denotes p<0.001 compared to RV-infected group at corresponding time point. (F-G) RV-1b infected mice were treated with vehicle or DP2 antagonist (OC000459) or DP1 antagonist (MK0524) or both DP2 and DP1 antagonist daily from day 99 and euthanized at 7 dpi. (F) Concentrations of IFN-γ in the BALF and TNF-α in the lung. (G) RV-1b infected mice were treated with vehicle, DP2 antagonist, fluticasone, or both DP2 antagonist and fluticasone daily from day 99 and euthanized at 7 dpi. Concentrations of IFN-γ in the BALF and TNF-α in the lung. Data are presented as box-and-whisker plots and are representative of two independent experiments showing similar results (n = 5-7 mice per group). * denotes p<0.05 and ** denotes p<0.01 compared to vehicle group. δ denotes p<0.05, δδ denotes p<0.01 and δδδ denotes p<0.001 compared to DP2 antagonist treated group.

### DP2 antagonism or DP1 agonism enhances NK cell IFN-γ production and alters the phenotype of alveolar macrophages

To identify the cell types producing IFN-γ and TNF-α at 7 dpi, we performed intracellular cytokine staining. RV inoculation decreased the number of IFN-γ producing cells, and this fall was arrested by DP2 antagonist or DP1 agonist treatment (Figure 5A). Several cell types expressed IFN-γ, however the predominant sources were CD4^+^ T cells, CD8^+^ T cells and NK cells (Figure 5B and Figure S5A), with only the number of IFN-γ-producing NK cells being increased by both treatments. TNF-α-producing cells increased in number following RV infection and were further increased by either intervention (Figure 5C). The predominant cell types expressing TNF-α were CD4^+^ T cells, type-2 classical dendritic cells (cDC2) and alveolar macrophages (AM); however, only TNF-α-producing AM numbers were increased by both treatments (Figure 5D-E). Consistent with other reports,^33^ viral inoculation significantly lowered lung AM numbers at 1 and 3 dpi. However, by 7 dpi, the AM population had recovered and was significantly elevated compared to the naïve controls (Figure 5F). As IFN-γ is known to contribute to the generation of CD86^+^CD206^−/low^ TNF-α-producing AMs,^34^ we next evaluated the effect of DP2 antagonism or DP1 agonism on the numbers of CD86^high^D206^−/low^ (M1-like macrophages) and CD86^−/low^CD206^high^ (M2-like macrophages) AMs during the RV exacerbation. As expected, CD86^−/low^CD206^high^ AMs were significantly elevated in mice with CEA (Figure 5G). Although neither intervention affected the total numbers of AMs (Figure 5F), treatment with either the DP2 antagonist or DP1 agonist attenuated the increase in CD86^−/low^CD206^high^ AMs between 3 and 7 dpi (Figure 5G), and increased CD86^high^CD206^−/low^ AM numbers (Figure 5G). A similar pattern was apparent when interstitial macrophages and monocytes were analyzed, with both populations showing fewer CD86^−/low^CD206^high^ AMs, particularly at 3 dpi (Figure S5C-D). Collectively, these data demonstrate that treatment with a DP2 antagonist or DP1 agonist during an RV-induced exacerbation increases IFN-γ-producing NK and T cell numbers, which is associated with a shift in the AM phenotype towards a CD86^High^CD206^−^ M1-like phenotype that produces TNF-α.

**FIGURE 5.**
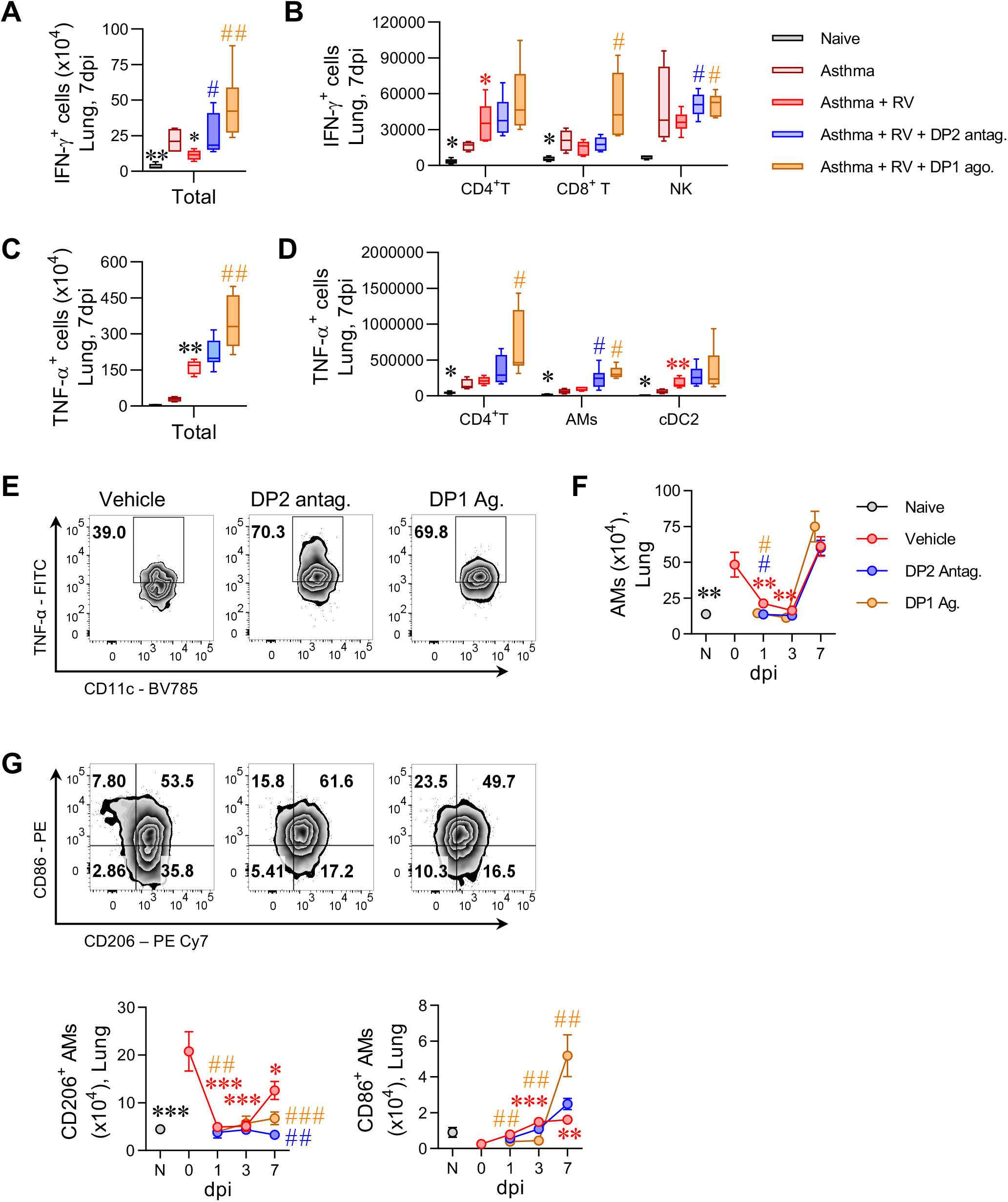
DP2 antagonism or DP1 agonism enhances NK cell IFN-γ production and alters the phenotype of alveolar macrophages. Mice with CEA were inoculated with RV-1b and treated with a DP2 antagonist or a DP1 agonist daily starting from day 99. Mice were euthanized at 1, 3 and 7 dpi. (A) Number of IFN-γ expressing cells in the lungs at 7 dpi. (B) Number of IFN-γ expressing CD4^+^ T, CD8^+^ T and NK cells in the lungs at 7 dpi. (C) Number of TNF-α expressing cells in the lungs at 7 dpi. (D) Number of TNF-α expressing CD4^+^ T, alveolar macrophages (AMs) and conventional type-2 dendritic cells (cDC2s) in the lungs at 7 dpi. (E) Representative flow cytometry plots showing TNF-α expressing AMs in the lungs. (F) Number of total AMs in the lungs. (G) Representative flow cytometry plots showing CD86 and CD206 expression on AMs and the total number of CD206-expressing AMs and CD86-expressing AMs. Data are presented as mean ± SEM or box-and-whisker plots and are pooled data from two independent experiments showing similar results (n = 4-8 mice per group). Statistical significance between different time points or different groups was determined using one-way ANOVA with Dunnett’s multiple comparison test. * denotes p<0.05; ** denotes p<0.01 and *** denotes p<0.001 compared to vehicle group at 0 dpi. # denotes p<0.05, ## denotes p<0.01 and ### denotes p<0.001 compared to RV-infected group at corresponding time point.

### DP2 antagonism mediates airway remodelling resolution via enhanced IFN-γ and TNF-α

To investigate whether IFN-γ contributes to the resolution of airway remodelling in response to DP2 antagonism, mice were treated with anti-IFN-γ and the DP2 antagonist during an RV-triggered exacerbation (Figure S6A). Anti-IFN-γ ablated the increase in IFN-γ levels induced by DP2 antagonism (Figure S6B), counteracted the resolving effects of DP2 antagonism on airway remodelling (Figure 6A-C) and ablated the decrease in TGF-β expression (Figure 6D and Figure S6C). Neutralization of IFN-γ reversed the suppressive effect of DP2 antagonism on type-2 inflammation and lowered PGD2 production (Figure S6D-G). Consistent with a role for IFN-γ in inducing CD86^+^ AMs, anti-IFN-γ attenuated the DP2 antagonist-mediated shift from CD206^+^ AMs to CD86^+^ AMs, and prevented the associated increase in TNF-α levels (Figure 6E-G). Taken together, these data suggest that the beneficial effects of DP2 antagonism on airway remodelling depend on the production of IFN-γ, which suppresses type-2 inflammation and the production of TGF-β1.

**Figure 6.**
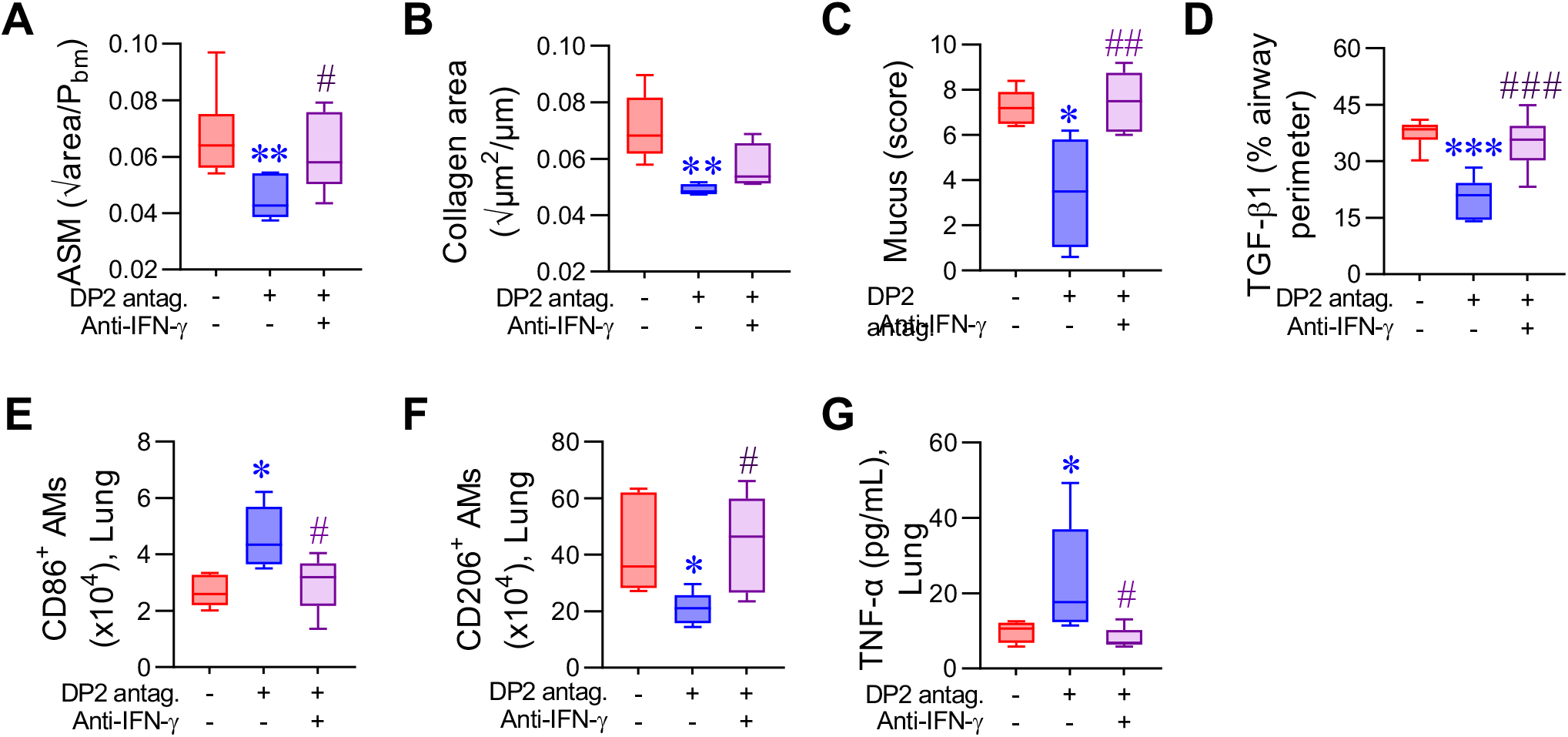
DP2 antagonism mediates airway remodelling resolution via enhanced IFN-γ. Mice with CEA were inoculated with RV-1b mice and treated with vehicle or DP2 antagonist +/− anti-IFN-γ. Mice were euthanized at 7 dpi. Quantification of (A) ASM area and (B) collagen deposition, (C) Mucus score and (D) TGF-β1 expression. (E-F) Number of CD86-expressing AM and CD206-expressing AMs in the lung. (G) Lung TNF-α levels. Data are presented as box-and-whisker plots and are representative of two independent experiments showing similar results (n = 5-10 mice per group). Statistical significance between different groups was determined using one-way ANOVA with Dunnett’s multiple comparison test. * denotes p<0.05; ** denotes p<0.01 and *** denotes p<0.001 compared to vehicle group. # denotes p<0.05, ## denotes p<0.01 and ### denotes p<0.001 compared to DP2 antagonist-treated group.

## DISCUSSION

In a mouse model of CEA, we demonstrated that DP2 antagonism during a RV-triggered exacerbation lowers established ASM mass and collagen deposition, mirroring the observation in the Phase II trial with fevipiprant. This phenotype was replicated by a DP1 agonist, and the beneficial effect of DP2 antagonism was ablated by DP1 antagonism, implicating the critical and beneficial role for endogenous PGD2-mediated activation of DP1. Corticosteroids dampened the RV-induced increase in endogenous PGD2 levels necessary for the resolution of ASM remodelling, and hence dual steroid/DP2-antagonist therapy impeded the effectiveness of DP2 antagonism. Mechanistically, we identified that DP2 antagonism lowers the expression of TGF-β1, a known ASM mitogen and pro-fibrotic cytokine, and that this effect was mediated by increased levels of IFN-γ.

In many of the Phase II trials of DP antagonists (fevipiprant, setipiprant, timapiprant), encouraging outcomes were observed.^9–15^ However, in the Phase III trials, fevipiprant led to only a modest reduction in the rates of asthma exacerbations in patients with inadequately controlled severe asthma, and thus failed to meet the primary efficacy endpoint.^16,17^ The underlying reasons for these disappointing results remain unclear, although the lower than expected number of exacerbations in the placebo arm decreased the window for improvement in the LUSTER studies.^16^ The 22% reduction in exacerbations in the patients that received the 450 mg dose suggests that a sub-group of patients were responsive.^16^ Potentially, the identification and targeting of the right study population, perhaps through measuring systemic or respiratory PGD2 production at baseline, or via stimulation of peripheral blood mononuclear cells *ex vivo*, might have improved the trial outcome. Importantly, the patients treated with fevipiprant or placebo were maintained on their standard of care asthma therapy, and therefore all the LUSTER trial participants were receiving inhaled corticosteroids and ∼10% of these patients were taking oral corticosteroids.^16^ Our study highlights that the effectiveness of DP2 antagonism is critically dependent on the production of endogenous PGD2, and thus the inhibitory effects of steroids on COX2 activity and expression may have counteracted the positive effects of DP2 antagonism in some patients, potentially explaining the mixed success of the various Phase II and III trials.^9–18,35^ Our findings revealed that DP2 antagonism in the absence of a RV-induced exacerbation has no benefit and that dual DP2 antagonism/steroid therapy is ineffective. Paradoxically, these observations suggest that therapy with a DP2 antagonist would work best in those patients who frequently exacerbate (given the need for endogenous PGD2), who are poorly compliant with their ICS/OCS therapy, or whose steroid insensitivity/resistance fails to dampen the production of PGD2. In a Phase II study that employed an experimental RV challenge of patients with asthma, DP2 antagonism did not affect exacerbation severity and of note, the investigators reported a negative correlation between PGD2 levels during RV infection and prescribed ICS dose.^35^ This observation and our findings would suggest that combining a DP2 antagonist with a DP1 agonist or a drug that induces endogenous PGD2, such as a TLR7 ligand, would elicit a more favourale outcome.

Consistent with the findings from the Phase II trial in asthma and our previous publication in the context of viral bronchiolitis,^11,19^ we observed that DP2 antagonism or DP1 agonism reversed established ASM area, mucus hypersecretion, and collagen deposition. Of note, we identified that inoculation with RV increased TGF-β1 expression in the ASM bundles, and that DP2 antagonism not only prevented this effect, but markedly decreased TGF-β1 expression by ASM cells. In contrast, fluticasone, despite lowering type-2 inflammation, had no effect on TGF-β1 expression and did not lower ASM area or collagen deposition. Consistent with previous reports,^19^ we observed that DP2 antagonism downregulated type-2 inflammation and induced a reciprocal upregulation of type-1 cytokines (IFN-γ and TNF-α), and moreover, we identified that the increase in IFN-γ expression mediated the pro-resolving effect of DP2 antagonism on ASM remodelling. Notably, corticosteroids also ablated the increase in IFN-γ and TNF-α expression induced by DP2 antagonism, consistent with their lack of effect on ASM area. Interestingly, NK cells were the only IFN-γ-producing cell type whose numbers were increased by both DP2 antagonism or DP1 agonism, although a limitation of our study is that we did not directly explore the role of this cell in mediating the protective effects of DP2 antagonism. The precise mechanism of action by which IFN-γ promoted the resolution of airway remodelling also remains an open question. IFN-γ can directly inhibit epidermal growth factor (EGF) and thrombin-induced ASM proliferation,^36^ and promotes the differentiation of CD86^high^CD206^−/low^ AMs,^34,37^ which in direct contrast to CD86^−/low^CD206^high^ AMs, inhibit fibrosis by releasing anti-fibrogenic factors like TNF-α.^38–40^ In our study, DP2 antagonism increased CD86^high^CD206^−/low^ AMs and decreased CD86^−/low^CD206^high^ AMs, and elevated TNF-α levels, and these phenotypes, all associated with a reduction in ASM area, were reversed following IFN-γ blockade. Further studies are needed to unravel the specific contribution of NK cells and AMs in mediating the resolution of airway remodelling.

In conclusion, using a high-fidelity preclinical model of CEA, we found that DP2 antagonism during a RV-induced asthma exacerbation resolves established ASM remodelling. Mechanistically, the beneficial effect of DP2 antagonism was mediated by increased levels of IFN-γ, which decreased TGF-β1 expression by ASM cells. Notably, the therapeutic efficacy of DP2 antagonism was dependent on endogenous PGD2-mediated activation of DP1, and ablated by dual therapy with a corticosteroid. The use of DP2 antagonists for the treatment of other indications that are characterized by small airway fibrosis and ASM remodelling, and for which steroids are not the mainstay treatment, such as chronic obstructive pulmonary disease, should be reevaluated.

## ACKNOWLEDGEMENTS

This work was supported by National Health and Medical Research Council (NHMRC, Australia) grants awarded to S.P. S.R. was funded by the doctorate program MOLIN (FWF, W1241) and an EMBO Short-Term Fellowship (8109). We thank all the staff from the QIMR Berghofer animal facility, flow cytometry laboratory and histology facility for their assistance, and Christina Kulis (Institute for Molecular Bioscience, University of Queensland) for the supply of the h-PGDS inhibitor, PK007.

## Abbreviations

ASM: airway smooth muscle
DP2: PGD2 receptor 2
PGD2: prostaglandin D2
TGF-β1: transforming growth factor β1

## AUTHORS CONTRIBUTIONS

M.A.U. and S.P. conceived the project, designed the experiments, analyzed and interpreted the data and wrote the manuscript. M.A.U., S.R., J.L. and B.F.C. conducted experiments, analyzed and interpreted the data. M.M.R., M.A.A.S., R.R., N.C., M.L. and L.B. performed experiments and analyzed the data. M.L.S. provided critical reagents and intellectual input. All the authors reviewed and approved the manuscript.

---

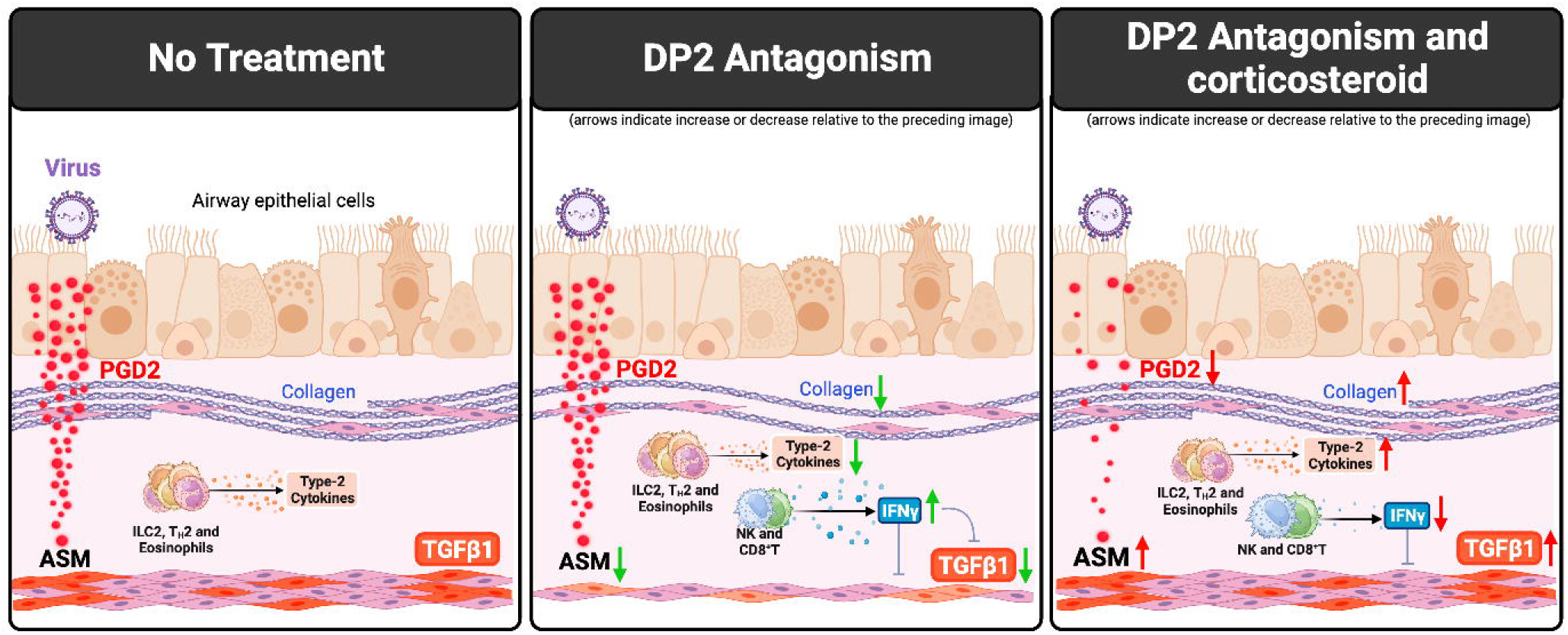

## REFERENCES

1. Busse WW, Lemanske RF, Jr., Gern JE. Role of viral respiratory infections in asthma and asthma exacerbations. Lancet. 2010;376(9743):826–834.

2. Jones RL, Noble PB, Elliot JG, et al. Airflow obstruction is associated with increased smooth muscle extracellular matrix. Eur Respir J. 2016;47(6):1855–1857.

3. Mauad T, Bel EH, Sterk PJ. Asthma therapy and airway remodeling. J Allergy Clin Immunol. 2007;120(5):997–1009; quiz 1010-1001.

4. Varricchi G, Ferri S, Pepys J, et al. Biologics and airway remodeling in severe asthma. Allergy. 2022;77(12):3538–3552.

5. Donovan GM, Langton D, Noble PB. Phenotype- and patient-specific modelling in asthma: Bronchial thermoplasty and uncertainty quantification. J Theor Biol. 2020;501:110337.

6. Domingo C, Palomares O, Sandham DA, Erpenbeck VJ, Altman P. The prostaglandin D(2) receptor 2 pathway in asthma: a key player in airway inflammation. Respir Res. 2018;19(1):189.

7. Johnston SL, Freezer NJ, Ritter W, O’Toole S, Howarth PH. Prostaglandin D2-induced bronchoconstriction is mediated only in part by the thromboxane prostanoid receptor. Eur Respir J. 1995;8(3):411–415.

8. Fajt ML, Gelhaus SL, Freeman B, et al. Prostaglandin D₂ pathway upregulation: relation to asthma severity, control, and TH2 inflammation. J Allergy Clin Immunol. 2013;131(6):1504–1512.

9. Diamant Z, Sidharta PN, Singh D, et al. Setipiprant, a selective CRTH2 antagonist, reduces allergen-induced airway responses in allergic asthmatics. Clin Exp Allergy. 2014;44(8):1044–1052.

10. Barnes N, Pavord I, Chuchalin A, et al. A randomized, double-blind, placebo-controlled study of the CRTH2 antagonist OC000459 in moderate persistent asthma. Clin Exp Allergy. 2012;42(1):38–48.

11. Saunders R, Kaul H, Berair R, et al. DP(2) antagonism reduces airway smooth muscle mass in asthma by decreasing eosinophilia and myofibroblast recruitment. Science translational medicine. 2019;11(479).

12. Gonem S, Berair R, Singapuri A, et al. Fevipiprant, a prostaglandin D2 receptor 2 antagonist, in patients with persistent eosinophilic asthma: a single-centre, randomised, double-blind, parallel-group, placebo-controlled trial. Lancet Respir Med. 2016;4(9):699–707.

13. Bateman ED, Guerreros AG, Brockhaus F, et al. Fevipiprant, an oral prostaglandin DP(2) receptor (CRTh2) antagonist, in allergic asthma uncontrolled on low-dose inhaled corticosteroids. Eur Respir J. 2017;50(2).

14. Erpenbeck VJ, Popov TA, Miller D, et al. The oral CRTh2 antagonist QAW039 (fevipiprant): A phase II study in uncontrolled allergic asthma. Pulm Pharmacol Ther. 2016;39:54–63.

15. Asano K, Sagara H, Ichinose M, et al. A Phase 2a Study of DP(2) Antagonist GB001 for Asthma. J Allergy Clin Immunol Pract. 2020;8(4):1275–1283.e1271.

16. Brightling CE, Gaga M, Inoue H, et al. Effectiveness of fevipiprant in reducing exacerbations in patients with severe asthma (LUSTER-1 and LUSTER-2): two phase 3 randomised controlled trials. Lancet Respir Med. 2021;9(1):43–56.

17. Castro M, Kerwin E, Miller D, et al. Efficacy and safety of fevipiprant in patients with uncontrolled asthma: Two replicate, phase 3, randomised, double-blind, placebo-controlled trials (ZEAL-1 and ZEAL-2). EClinicalMedicine. 2021;35:100847.

18. Moss MH, Lugogo NL, Castro M, et al. Results of a Phase 2b Trial With GB001, a Prostaglandin D(2) Receptor 2 Antagonist, in Moderate to Severe Eosinophilic Asthma. Chest. 2022;162(2):297–308.

19. Werder RB, Lynch JP, Simpson JC, et al. PGD2/DP2 receptor activation promotes severe viral bronchiolitis by suppressing IFN-lambda production. Science translational medicine. 2018;10(440).

20. Lynch JP, Werder RB, Simpson J, et al. Aeroallergen-induced IL-33 predisposes to respiratory virus-induced asthma by dampening antiviral immunity. J Allergy Clin Immunol. 2016;138(5):1326–1337.

21. Werder RB, Ullah MA, Rahman MM, et al. Targeting the P2Y(13) Receptor Suppresses IL-33 and HMGB1 Release and Ameliorates Experimental Asthma. Am J Respir Crit Care Med. 2022;205(3):300–312.

22. Werder RB, Zhang V, Lynch JP, et al. Chronic IL-33 expression predisposes to virus-induced asthma exacerbations by increasing type 2 inflammation and dampening antiviral immunity. J Allergy Clin Immunol. 2018;141(5):1607–1619.e1609.

23. Curren B, Ahmed T, Howard DR, et al. IL-33-induced neutrophilic inflammation and NETosis underlie rhinovirus-triggered exacerbations of asthma. Mucosal Immunol. 2023.

24. Holt PG, Sly PD. Viral infections and atopy in asthma pathogenesis: new rationales for asthma prevention and treatment. Nat Med. 2012;18(5):726–735.

25. Ullah MA, Rittchen S, Li J, Hasnain SZ, Phipps S. DP1 prostanoid receptor activation increases the severity of an acute lower respiratory viral infection in mice via TNF-α-induced immunopathology. Mucosal Immunol. 2021;14(4):963–972.

26. Vijay R, Fehr AR, Janowski AM, et al. Virus-induced inflammasome activation is suppressed by prostaglandin D(2)/DP1 signaling. Proc Natl Acad Sci U S A. 2017;114(27):E5444–e5453.

27. Boulet LP, Turcotte H, Laviolette M, et al. Airway hyperresponsiveness, inflammation, and subepithelial collagen deposition in recently diagnosed versus long-standing mild asthma. Influence of inhaled corticosteroids. AmJRespirCrit Care Med. 2000;162(4 Pt 1):1308–1313.

28. Chakir J, Shannon J, Molet S, et al. Airway remodeling-associated mediators in moderate to severe asthma: effect of steroids on TGF-beta, IL-11, IL-17, and type I and type III collagen expression. J Allergy Clin Immunol. 2003;111(6):1293–1298.

29. Ojiaku CA, Yoo EJ, Panettieri RA, Jr. Transforming Growth Factor β1 Function in Airway Remodeling and Hyperresponsiveness. The Missing Link? Am J Respir Cell Mol Biol. 2017;56(4):432–442.

30. Flood-Page P, Menzies-Gow A, Phipps S, et al. Anti-IL-5 treatment reduces deposition of ECM proteins in the bronchial subepithelial basement membrane of mild atopic asthmatics. J Clin Invest. 2003;112(7):1029–1036.

31. Redington AE, Meng QH, Springall DR, et al. Increased expression of inducible nitric oxide synthase and cyclo-oxygenase-2 in the airway epithelium of asthmatic subjects and regulation by corticosteroid treatment. Thorax. 2001;56(5):351–357.

32. Aksoy MO, Li X, Borenstein M, Yi Y, Kelsen SG. Effects of topical corticosteroids on inflammatory mediator-induced eicosanoid release by human airway epithelial cells. J Allergy Clin Immunol. 1999;103(6):1081–1091.

33. Califano D, Furuya Y, Metzger DW. Effects of Influenza on Alveolar Macrophage Viability Are Dependent on Mouse Genetic Strain. J Immunol. 2018;201(1):134–144.

34. Saradna A, Do DC, Kumar S, Fu QL, Gao P. Macrophage polarization and allergic asthma. Transl Res. 2018;191:1–14.

35. Farne H, Glanville N, Johnson N, et al. Effect of CRTH2 antagonism on the response to experimental rhinovirus infection in asthma: a pilot randomised controlled trial. Thorax. 2022;77(10):950–959.

36. Amrani Y, Tliba O, Choubey D, et al. IFN-gamma inhibits human airway smooth muscle cell proliferation by modulating the E2F-1/Rb pathway. Am J Physiol Lung Cell Mol Physiol. 2003;284(6):L1063–1071.

37. Guilliams M, van de Laar L. A Hitchhiker’s Guide to Myeloid Cell Subsets: Practical Implementation of a Novel Mononuclear Phagocyte Classification System. Front Immunol. 2015;6:406.

38. Frankel SK, Cosgrove GP, Cha SI, et al. TNF-alpha sensitizes normal and fibrotic human lung fibroblasts to Fas-induced apoptosis. Am J Respir Cell Mol Biol. 2006;34(3):293–304.

39. Yamane K, Ihn H, Asano Y, Jinnin M, Tamaki K. Antagonistic effects of TNF-alpha on TGF-beta signaling through down-regulation of TGF-beta receptor type II in human dermal fibroblasts. J Immunol. 2003;171(7):3855–3862.

40. Wynes MW, Edelman BL, Kostyk AG, et al. Increased cell surface Fas expression is necessary and sufficient to sensitize lung fibroblasts to Fas ligation-induced apoptosis: implications for fibroblast accumulation in idiopathic pulmonary fibrosis. J Immunol. 2011;187(1):527–537.

