## Supplementary figures and images for "Dual therapy with corticosteroid ablates the beneficial effect of DP2 antagonism in chronic experimental asthma"

### Supplemental Figures

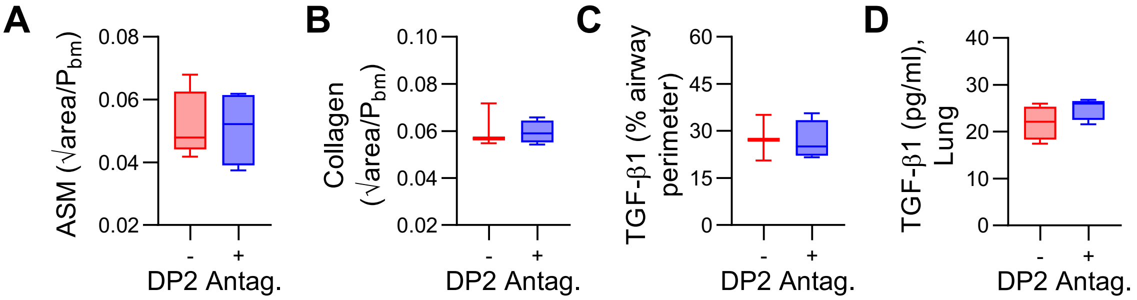

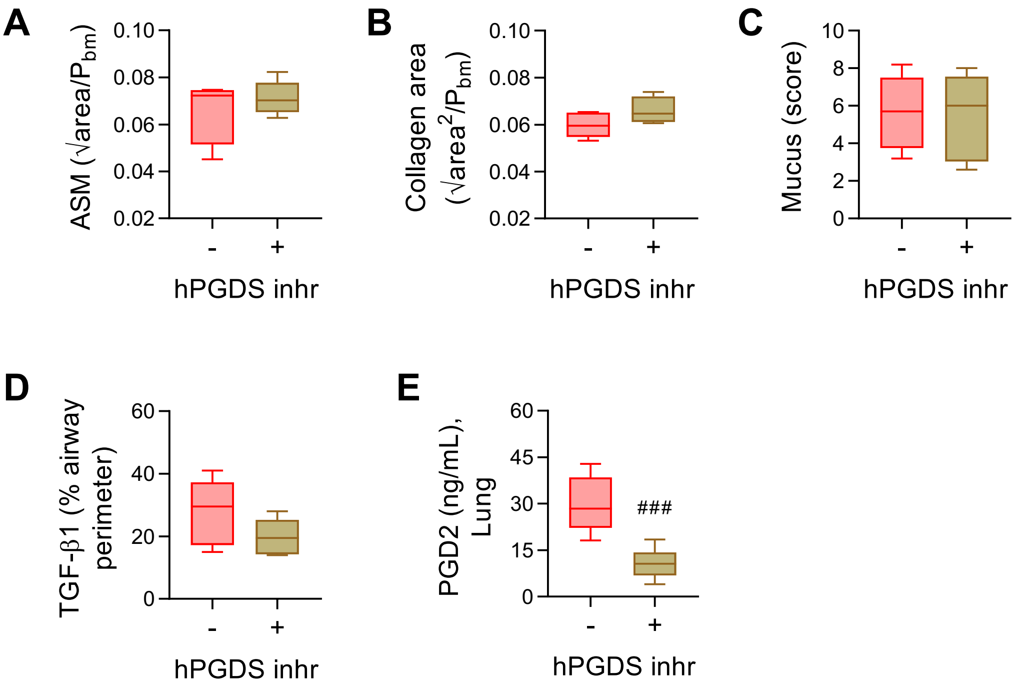

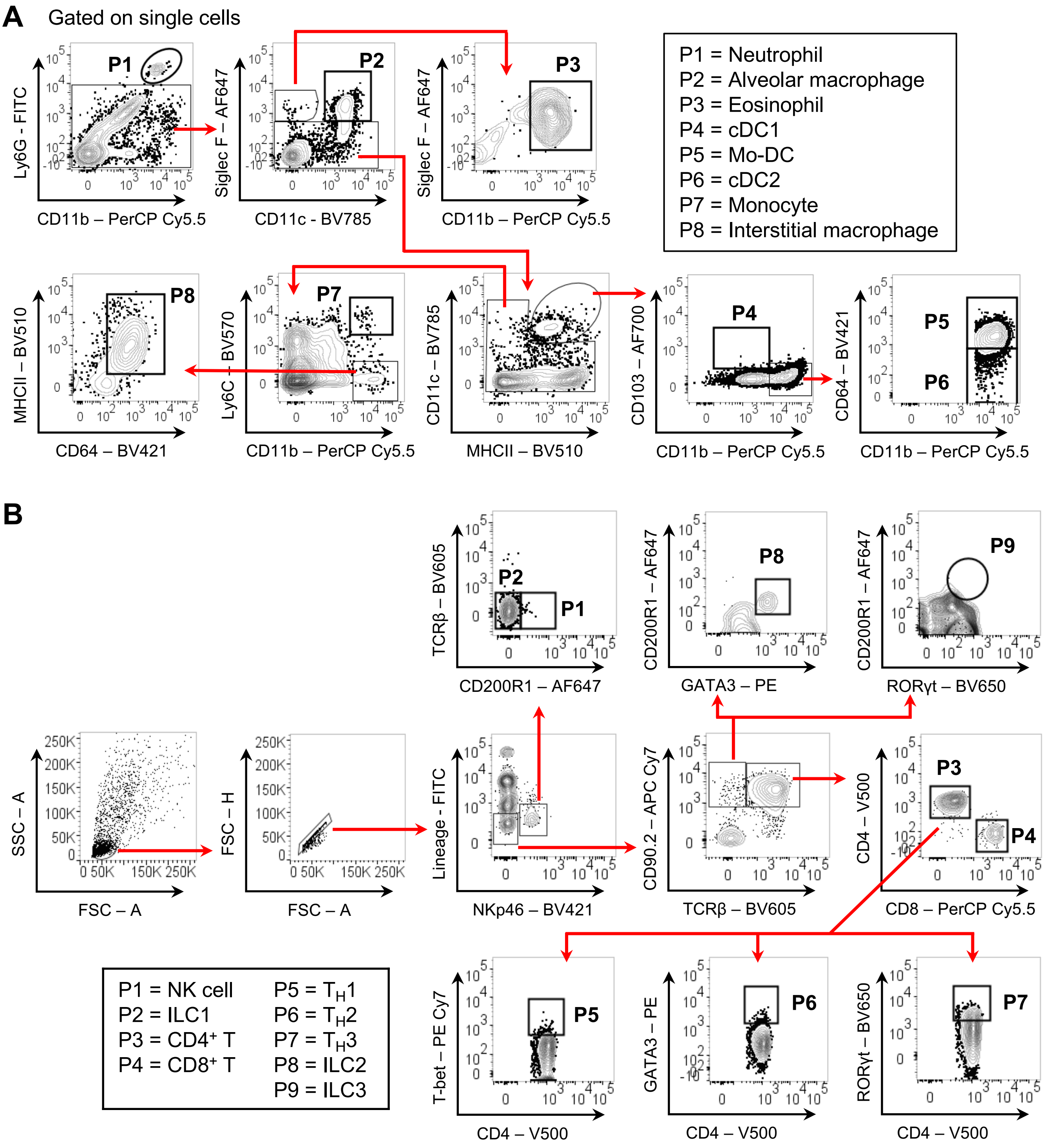

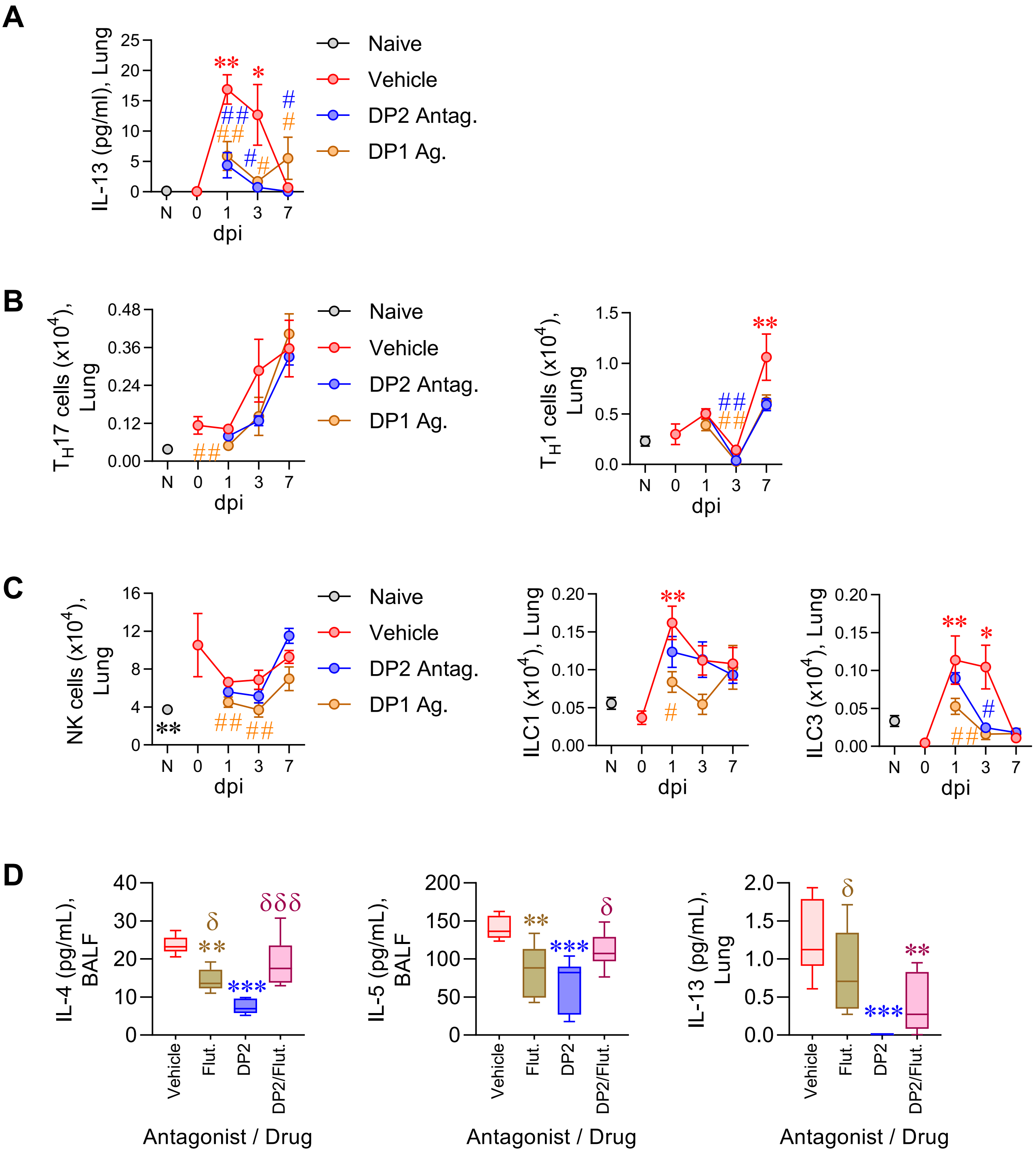

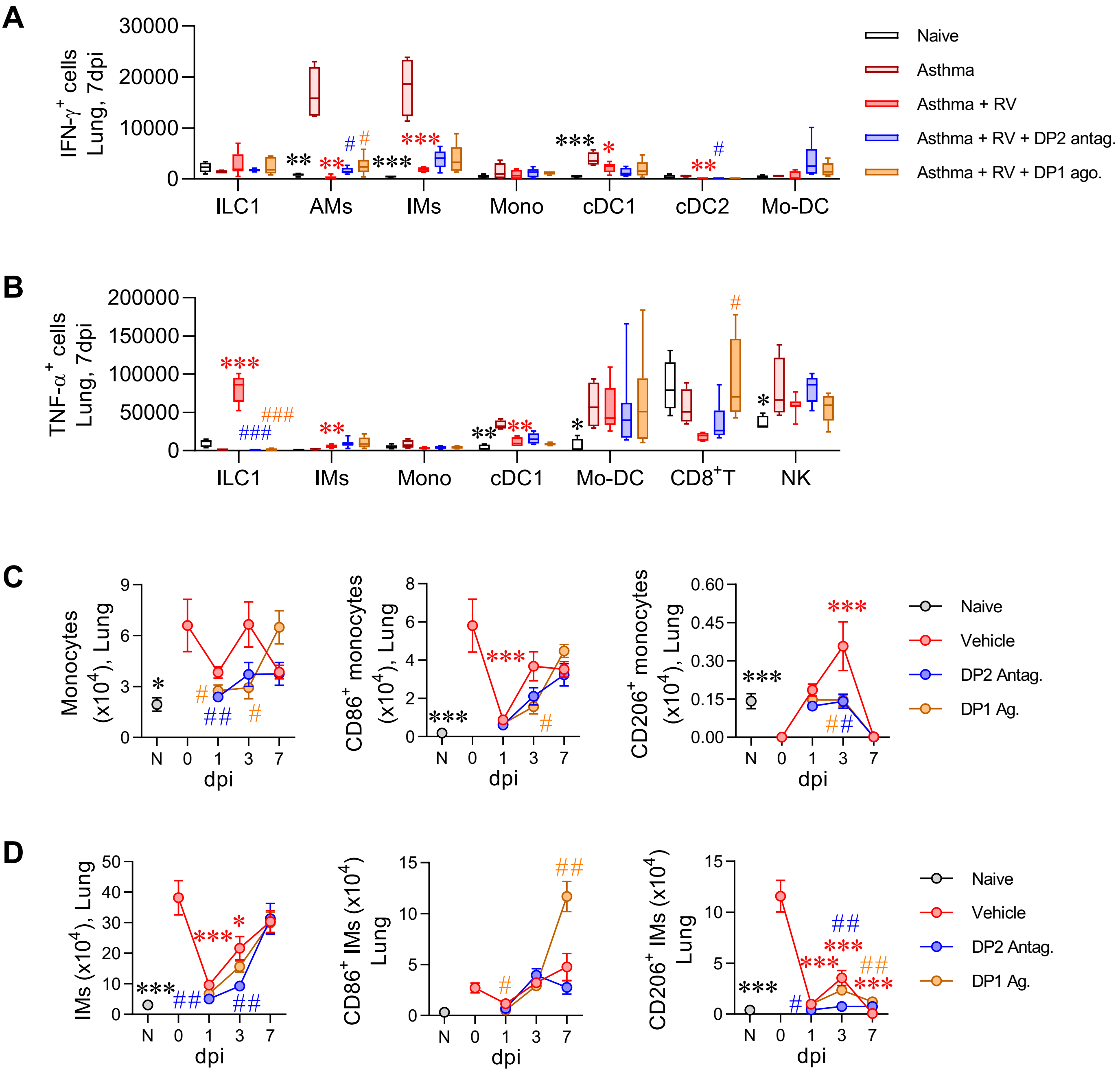

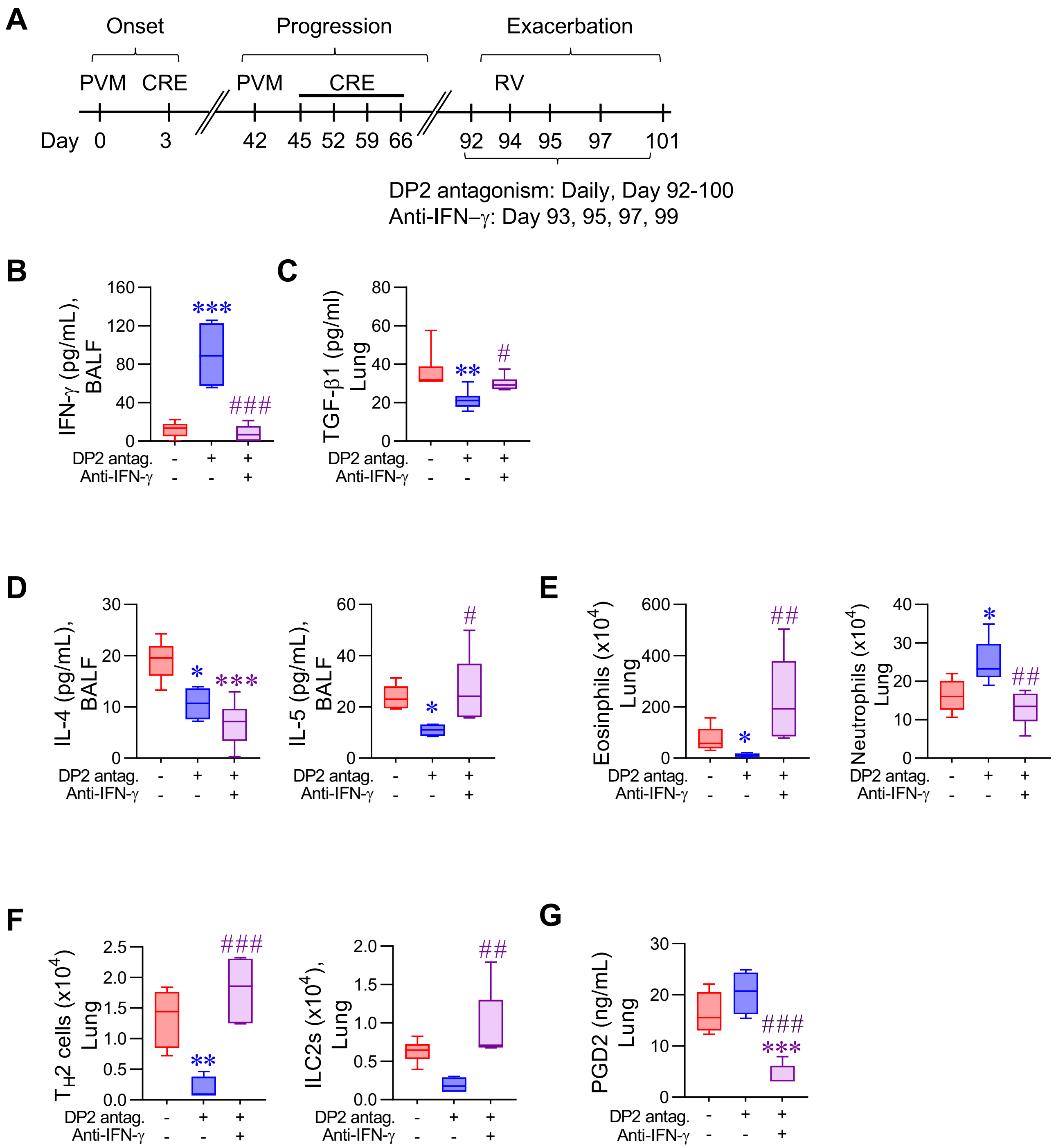
