## Supplemental Figure Legends for "Dual therapy with corticosteroid ablates the beneficial effect of DP2 antagonism in chronic experimental asthma"

### SUPPLEMENTARY FIGURE LEGENDS

#### FIGURE S1. DP2 antagonism does not ameliorate airway remodelling in mice with chronic asthma.

Mice with chronic asthma were treated with DP2 antagonist, OC000459 daily from day 99 and were euthanized on day 108. (A) ASM area. (B) Collagen area. (C) Quantification of TGF-β1 immunoreactivity around airways. (D) Lung TGF-β1 levels in homogenates. Data are presented as box-and-whisker plots and are representative of two independent experiments showing similar results (n = 3-7 mice per group). Group differences were analyzed using Mann-Whitney U test.

#### FIGURE S2. Inhibition of h-PGDS does not resolve airway remodelling during RV-induced exacerbation of chronic asthma.

Mice with chronic asthma were inoculated with RV-1b on day 101. Separate group of RV-1b infected mice were treated with h-PGDS inhibitor (PK007) daily from day 99 onwards and euthanized 3 days post infection. (A) ASM area. (B) Collagen area. (C) Muc5ac score. (D) Quantification of TGF-β1 immunoreactivity around airways. (E) Lung PGD2 levels. Data are presented as box-and-whisker plots and are representative of two independent experiments showing similar results (n = 4-9 mice per group). Group differences were analyzed using Mann-Whitney U test. ### denotes p<0.001.

#### FIGURE S3. Gating strategies of immune cells in the lung.

Representative flow cytometry plots depicting the gating strategies for lung immune cells.

#### FIGURE S4. Relates to Figure 4.

Mice with chronic asthma were inoculated with RV-1b and treated with DP2 antagonist or DP1 agonist or vehicle daily starting from day 99. Mice were euthanized at 1, 3 and 7 dpi. (A) IL-13 expression in the lungs. (B) Number of T_H_17 (RORγt^+^CD4^+^ T) and T_H_1 (T-bet^+^CD4^+^ T) cells in the lungs. (C) Number of NK cells (CD3ε^−^CD19^−^NK1.1^+^NKp46^+^CD200R1^-^), ILC1s (CD3ε^−^CD19^−^CD45R^−^CD11c^−^Gr‐1^−^ NK1.1^+^NKp46^+^CD90.2^+^CD200R1^+^T-bet^+^) and ILC3s (CD3ε^−^CD19^−^CD45R^−^CD11c^−^ Gr‐1^−^CD11b^−^ CD90.2^+^CD200R1^+^RORγt^+^) in the lungs. Data are presented as mean ± SEM and are representative of two independent experiments showing similar results (n = 4-7 mice per group). * denotes p<0.05 and ** denotes p<0.01 compared to Vehicle group. # denotes p<0.05 and ## denotes p<0.01 compared to RV-infected group at corresponding time point. (D) RV-1b infected mice were treated with vehicle or DP2 antagonistor fluticasone or both DP2 antagonist and fluticasone daily from day 99 and euthanized at 7 dpi. Concentrations of IL-4 and IL-5 in the BALF and IL-13 in the lungs. Data are presented as box-and-whisker plots and are representative of two independent experiments showing similar results (n = 4-7 mice per group). Statistical significance between different time points or different groups was determined using one-way ANOVA with Dunnett’s multiple comparison test. * denotes p<0.05; ** denotes p<0.01 and *** denotes p<0.001 compared to vehicle group. δ denotes p<0.05 and δδδ denotes p<0.001 compared to DP2 antagonist (OC000459) treated group.

#### FIGURE S5. Relates to Figure 5.

Mice with chronic asthma were inoculated with RV-1b and treated with a DP2 antagonist or a DP1 agonist daily starting from day 99. Mice were euthanized at 1, 3 and 7 dpi. (A) Number of IFN-γ expressing ILC1, AMs, interstitial macrophages (IMs), monocytes (Mono), conventional type-1 dendritic cells (cDC1), conventional type-2 dendritic cells (cDC2) and monocyte-derived dendritic cells (Mo-DC) in the lungs at 7 dpi. (B) Number of TNF-α expressing ILC1, IMs, Mono, cDC1, Mo-DC, CD8^+^T and NK cells in the lungs at 7 dpi. (C) Number of total monocytes (CD11b^+^Ly6C^+^SiglecF^-^) and CD86-expressing and CD206-expressing monocytes in the lungs. (D) Number of total interstitial macrophages (IM; SiglecF^-^Ly6C^-^CD11b^+^MHCII^+^CD64^+^) and CD86-expressing and CD206-expressing IMs in the lungs. Data are presented as mean ± SEM or box-and-whisker plots and are pooled data from two independent experiments showing similar results (n = 4-8 mice per group). Statistical significance between different time points or different groups was determined using one-way ANOVA with Dunnett’s multiple comparison test. * denotes p<0.05; ** denotes p<0.01 and *** denotes p<0.001 compared to asthma or vehicle group. # denotes p<0.05, ## denotes p<0.01 and ### denotes p<0.001 compared to RV-infected group at corresponding time point.

**Figure S6. Relates to Figure 6.**

(A) Study design. RV-1b infected mice were given vehicle or DP2 antagonist and treated with anti-IFN-γ. Mice were euthanized at 7 dpi. (B) IFN-γ expression in the BALF. (C) TGF-β1 levels in homogenates. (D) Concentrations of IL-4 and IL-5 in the BALF. (E) Number of eosinophils and neutrophils in the lungs. (F) Number of T_H_2 cells and ILC2s in the lungs. (G) Lung PGD2 levels. Data are presented as box-and-whisker plots and are representative of two independent experiments showing similar results (n = 4-6 mice per group). Statistical significance between different groups was determined using one-way ANOVA with Dunnett’s multiple comparison test. * denotes p<0.05; ** denotes p<0.01 and *** denotes p<0.001 compared to vehicle group. # denotes p<0.05, ## denotes p<0.01 and ### denotes p<0.001 compared to OC000459-treated group.
