## Supplementary Table S1 for "Dual therapy with corticosteroid ablates the beneficial effect of DP2 antagonism in chronic experimental asthma"

### SUPPLEMETARY TABLE

**Table S1**. Information of antibodies, kits and reagents used in this study

| **Name** | **Supplier** |
| --- | --- |
| ***Anti-mouse antibodies*** |  |
| B220-FITC (RA3-6B2) | Biolegend (CA, USA) |
| CD103-AF700 (2E7) | Biolegend (CA, USA) |
| CD11b-PerCp-Cy5.5, BV605 (M1/70) | BD Biosciences (CA, USA) |
| CD11c-BV785 (N418) | Biolegend (CA, USA) |
| CD19-FITC (1D3/CD19) | Biolegend (CA, USA) |
| CD200R1-AF647 (OX-110) | BD Biosciences (CA, USA) |
| CD206-PECy7 (C068C2) | Biolegend (CA, USA) |
| CD3ε-FITC (17A2) | Biolegend (CA, USA) |
| CD4-FITC (RM4-5) | Biolegend (CA, USA) |
| CD4-V500 (RM4-5) | BD Biosciences (CA, USA) |
| CD64-BV421 (X54-5/7.1) | Biolegend (CA, USA) |
| CD8-PerCp-Cy5.5 (53-6.7) | Biolegend (CA, USA) |
| CD80-APC/Fire750 (16-10A1) | Biolegend (CA, USA) |
| CD86-PE (GL-1) | Biolegend (CA, USA) |
| CD90.2-APC Cy7 (53-2.1) | BD Biosciences (CA, USA) |
| F4/80-FITC, APC, APC Cy7 (BM8) | Biolegend (CA, USA) |
| FoxP3-AF647 (MF23) | BD Biosciences (CA, USA) |
| GATA3-PE (TWAJ) | eBiosciences (CA, USA) |
| Gr-1-FITC (RB6-8C5) | Biolegend (CA, USA) |
| IFN-γ-PE Cy7 (XMG1.2) | Biolegend (CA, USA) |
| Ly6C-BV570 (HK1.4) | Biolegend (CA, USA) |
| Ly6G-FITC (1A8) | BD Biosciences (CA, USA) |
| MHCII-BV510 (M5/114.15.2) | Biolegend (CA, USA) |
| Muc5ac (45M1) | Invitrogen (CA, USA) |
| NKp46-BV421 (29A1.4) | Biolegend (CA, USA) |
| RORγt-BV650 (Q31-378) | BD Biosciences (CA, USA) |
| Siglec F-PE (E50-2440) | BD Biosciences (CA, USA) |
| T-bet-PE Cy7 (4B10) | Biolegend (CA, USA) |
| TCR-β-BV605 (H57-597) | Biolegend (CA, USA) |
| TER119-FITC (TER-119) | Biolegend (CA, USA) |
| TGF-β1 (polyclonal) | Abcam (USA) |
| TNF-α-FITC (MP6-XT22) | BD Biosciences (CA, USA) |
| α-Smooth muscle actin (1A4) | Sigma-Aldrich (Mo, USA) |
| ***Kits*** |  |
| FoxP3/TF fixation/permeabilization kit | eBiosciences (CA, USA) |
| Mouse IFN-γ ELISA Kit | Biolegend (CA, USA) |
| Mouse IL-13 enhanced sensitivity flex set | BD Biosciences (CA, USA) |
| Mouse IL-17A ELISA Kit | Biolegend (CA, USA) |
| Mouse IL-4 ELISA Kit | BD Biosciences (CA, USA) |
| Mouse IL-5 ELISA Kit | BD Biosciences (CA, USA) |
| Mouse TGF-β1 ELISA kit | R&D systems (MN, USA) |
| Mouse TNF-α enhanced sensitivity flex set | BD Biosciences (CA, USA) |
| PGD2-MOX Express ELISA kit | Cayman Chemicals (Michigan, USA) |
| Zombie Aqua™ Fixable Viability Kit | Biolegend (CA, USA) |
| ***Reagents*** |  |
| Brefeldin A | Biolegend (CA, USA) |
| BW245c | Cayman Chemical (MI, USA) |
| Cockroach extract | Greer Laboratories (NC, USA) |
| Hyclone fetal calf serum | Cytiva (USA) |
| Fluticasone | Sigma-Aldrich (MO, USA) |
| Glycerol gel | Sigma-Aldrich (MO, USA) |
| Hematoxylin, Dako | Agilent Technologies (Australia) |
| IGEPAL® CA-630 | Sigma-Aldrich (MO, USA) |
| Ionomycin | Sigma-Aldrich (MO, USA) |
| MK-0524 | Cayman Chemical (MI, USA) |
| OC000459 | Cayman Chemical (MI, USA) |
| Phorbol 12-myristate 13-acetate (PMA) | Sigma-Aldrich (MO, USA) |
| Sirius red | Sigma-Aldrich (MO, USA) |
| Sodium chloride | Sigma-Aldrich (MO, USA) |
| Sodium deoxycholate | Sigma-Aldrich (MO, USA) |
| Sodium dodecyl sulfate | Sigma-Aldrich (MO, USA) |
| Soluble IL-13Rα2 | Pfizer, NY, USA |
| Tris-base | Sigma-Aldrich (MO, USA) |
