## Supplementary material for "Dual therapy with corticosteroid ablates the beneficial effect of DP2 antagonism in chronic experimental asthma": Methods

### Lung tissue harvesting and processing

Following euthanasia, bronchoalveolar lavage (BAL) was performed. The cell-free supernatant was stored at -80^0^C prior to analysis. The left lobe was then immediately processed for flow cytometry analysis. The superior right lobe was placed in neutral buffered formalin for fixation and processed for histological analysis. The right inferior lobe was snap frozen in protein assay buffer containing sodium chloride (150 mM), IGEPAL^®^ CA-630 (1%), sodium deoxycholate (0.5%), sodium dodecyl sulfate (0.1%) and Tris buffer (50mM) and stored -80^0^C. This lobe was later homogenized with a tissue-tearor (Biospec, OK, USA), centrifuged at 13,000 rpm for 10 min, and the supernatant harvested and stored at -80^0^C for ELISA.

### Flow cytometry

Single-cell suspension was prepared from lung tissue as described previously.^1^ Red blood cells were lysed with Gey’s lysis buffer, and the cells re-suspended and washed in FASC buffer (PBS supplemented with 2% fetal calf serum, FCS). Cells were incubated with 2.4G2 antibody for 30 minutes at 4^0^C to prevent non-specific binding, then stained with the surface antibody cocktail for 30 minutes at 4^0^C. The list of antibodies used for flow cytometry is summarized in Table S1. To detect intracellular transcription factors, the cells were first stained with surface antibodies then permeabilized and fixed using FoxP3/transcription factor fixation/permeabilization kit (eBiosciences; CA, USA). After washing with permeabilization buffer, the cells were stained with fluorochrome-conjugated antibodies against ROR-γτ, GATA3 or Tbet. For intracellular cytokine staining, cells were stimulated with phorbol 12-myristate 13-acetate (PMA; 50 ng/ml) and ionomycin (1 µg/ml) in the presence of brefeldin A (20 µg/ml) at 37^0^C for 3 hours, then washed with FACS buffer and stained with surface antibodies as above. Cells were permeabilized and fixed using FoxP3/transcription factor fixation/permeabilization kit and stained with appropriate antibodies. The samples were acquired on a BD Fortessa IV flow cytometer (BD Biosciences, USA) using the FACSDiva software (version 8, BD Biosciences, USA). Data was analyzed using FlowJo software (Version 10.6; TeeStar, USA) and gating strategy has been depicted in Supplementary Figure S1.

### Histology

Paraffin-embedded lung sections (5 µm thick) were prepared and stained with the appropriate antibodies as described previously.^2,3^ Briefly, lung sections were deparaffinized and rehydrated. Then, the sections were immersed in citrate buffer and heated in a pressure cooker for antigen retrieval. Sections were permeabilized using 0.6% Tween-20 and then incubated with 10% goat serum in phosphate-buffered saline (PBS) for thirty minutes at room temperature. Subsequently, sections were incubated with the following primary antibodies (details are in supplementary Table S1): mouse anti-Muc5ac (1:400), mouse anti-α-Smooth muscle actin (α-SMA; 1:800) and rabbit anti-TGF-β1 (1:400). The following day, sections were washed and incubated with the appropriate alkaline-phosphatase secondary antibody. The color was developed with fast-red reagent. Sections were counter-stained with hematoxylin and mounted with glycerol gel. Tissue was stained with picrosirius red (PSR) to visualize collagen expression. All the stained slides were scanned using Aperio AT Turbo (Leica Biosystems, Wetzlar, Germany). Muc5ac staining was scored on a scale of 1 to 5 for the percentage of airway epithelial cells expressing Muc5ac and 1 to 5 for the percentage of the airway with Muc5ac plugging with a maximum score of 10.^4^ The sum of ASM or collagen area around the small airways was quantified using Aperio ImageScope software (Leica Biosystems, Wetzlar, Germany) and the square root of the ASM or collagen area was presented per micrometer of basement membrane circumference.^5^ TGF-β1 expression around the small airways was expressed as the fraction of airway perimeter with adjacent TGF-β1 expressing cells.^2^

### ELISA assay

The expression of cytokines, chemokines, PGD2 and dsDNA was measured using commercially available kits as per the manufacturer’s instructions (supplementary Table S1).

### Statistical analysis

All the experiments were performed at least two times. Where data are presented as box-and-whisker plots, the boxes represent quartiles and whiskers indicate the range. The software GraphPad Prism (version 8, USA) was used for statistical analysis. Group differences were analyzed by the Mann-Whitney U test or one-way ANOVA with Dunnett’s multiple comparison tests as appropriate. Significance was set at a P value less than 0.05 for all tests.
